# Association between the rifampicin resistance mutations and rifabutin susceptibility in *Mycobacterium tuberculosis*: a meta-analysis

**DOI:** 10.1101/2023.07.13.548878

**Authors:** Wenli Wang, Hongjuan Zhou, Long Cai, Tingting Yang

## Abstract

Some rifampicin-resistant *Mycobacterium tuberculosis* (MTB) strains were susceptible to rifabutin (RFB) and may be amenable to treatment with RFB. We performed a meta-analysis of available cross-sectional studies to determine which RIF-resistance mutations were associated with rifabutin susceptibility. We identified studies through PubMed, Web of Science, Embase, and Cochrane Library up to June 1, 2023. Studies that met our criteria were those that investigated *rpoB* mutations and reported phenotypic drug susceptibility for RIF and RFB. The relationship between RIF-resistance mutations to RFB-susceptibility was evaluated using odds ratio (OR). Twenty-five studies comprised 4,333 clinical RIF-resistant MTB isolates from 21 different countries met our criteria for inclusion. Of these isolates, 21.00% (910/4333) were susceptible to RFB. We found seven RIF-resistance mutations were high confidence (OR>10) in predicting RFB-susceptibility, which were D435V, D435Y, D435F, H445L, L430R, S441L, and S441Q. Among strains carrying these mutations, 83.01% (435/524) were susceptible to RFB. The minimum inhibition concentrations (MICs) of these strains revealed that they had low MIC (D435V, D435F, H445L, and D435Y) or were susceptible (S441L) for RFB and exhibited a significant lack of correlation between MICs to RIF and RFB. Mutations such as H445C, H445G, H445N, L430P, and L452P showed a moderate confidence (5<OR≤10) in prediction of RFB-susceptibility. Of these mutants, 62.16% (69/111) were susceptible to RFB. The most common RIF-resistance mutations S450L, as well as S450W, were associated with RFB-resistance (OR<1). These results provide a theoretical basis for molecular detection of RFB-susceptible TB and alternative treatment with RFB in MDR/RR-TB patients.

## Introduction

Tuberculosis (TB) remains one of the leading causes of death worldwide and the emergence and spread of drug resistance have posed significant challenges to its control (1). In 2021, there were 450,000 new cases of rifampicin-resistant tuberculosis (RR-TB) globally, a 3.1% increase from 437,000 in 2020 (1). Approximately 78% of RR-TB patients have multidrug-resistant tuberculosis (MDR-TB), being resistant to both rifampicin and isoniazid, the two most effective anti-tuberculosis drugs (2). MDR/RR greatly diminishes the likelihood of successful treatment in these TB patients, with a treatment success rate of 60% compared to 86% in drug-susceptible TB patients, and a high mortality rate of 16% (1). Despite the availability of newer drugs such as bedaquiline, delamanid, linezolid, and clofazimine for the treatment of MDR-TB, challenges such as high drug prices, treatment complexity, and significant adverse reactions persist (3).

Some MDR/RR-TB patients may be treated with rifabutin (RFB), which belongs to the same class of drugs as rifampicin (RIF). RFB has a treatment effect similar to that of rifampicin and has lower toxicity than second-line drugs (4, 5). Multiple studies have reported that some clinical *Mycobacterium tuberculosis* (MTB) strains with certain known RIF-resistance mutations were susceptible to rifabutin (6, 7). Clinical evidence from Jo et al. (8) showed favorable outcomes in some RR-TB patients treated with RFB. These indicated that the identification of RFB-susceptible strains within MDR/RR-MTB can aid in the precise clinical use of RFB in the treatment of the corresponding patients.

Although the known RIF-resistance mutations had been proven to have high accuracy in predicting RIF resistance, their relationship with RFB susceptibility remains unclear. Several studies had explored this relationship. Early studies correlated PCR-detected *rpoB* gene or RRDR mutations with the phenotypic drug susceptibility test (pDST) results of strains to RIF and RFB (9-12). In recent years, more methods have been applied to both pDST and mutation detection, resulting in a wealth of phenotype-genotype association data (13-16). However, due to sample size and mutation frequency limitations, these studies have not definitively established the relationship between RIF-resistance mutations and RFB-susceptibility (17-20). Whitfield et al. reported on the resistance levels to RIF and RFB in MTB strains carrying different *rpoB* mutations in a sample of 2,045 isolates in 2018, but they did not further assess the statistical significance of the association between mutations and RFB susceptibility (21). To systematically assess such relationships, we conducted a meta-analysis using the relevant phenotype-genotype associated data from existing studies.

## Results

### Literature search

A total of 327 articles were retrieved from the PubMed, Web of Science, Embase, and Cochrane Library databases (Table S1 and Fig. 1). Of these, 268 articles were removed as they were duplicates (n=120), irrelevant (n=145) and full-text inaccessible (n=3). Through reading the full text of the remaining 59 articles, we further excluded 34 articles that focused on experimental strains (n=3), did not provide detailed information on mutations and/or pDST results (n=20), were non-English (n=4), were case-control studies (n=4), or lacked the rifabutin-susceptible group (n=3). Finally, a total of 25 articles were included for meta-analysis. The review protocol was registered on the PROSPERO database (CRD42023431207).

**Fig 1.**
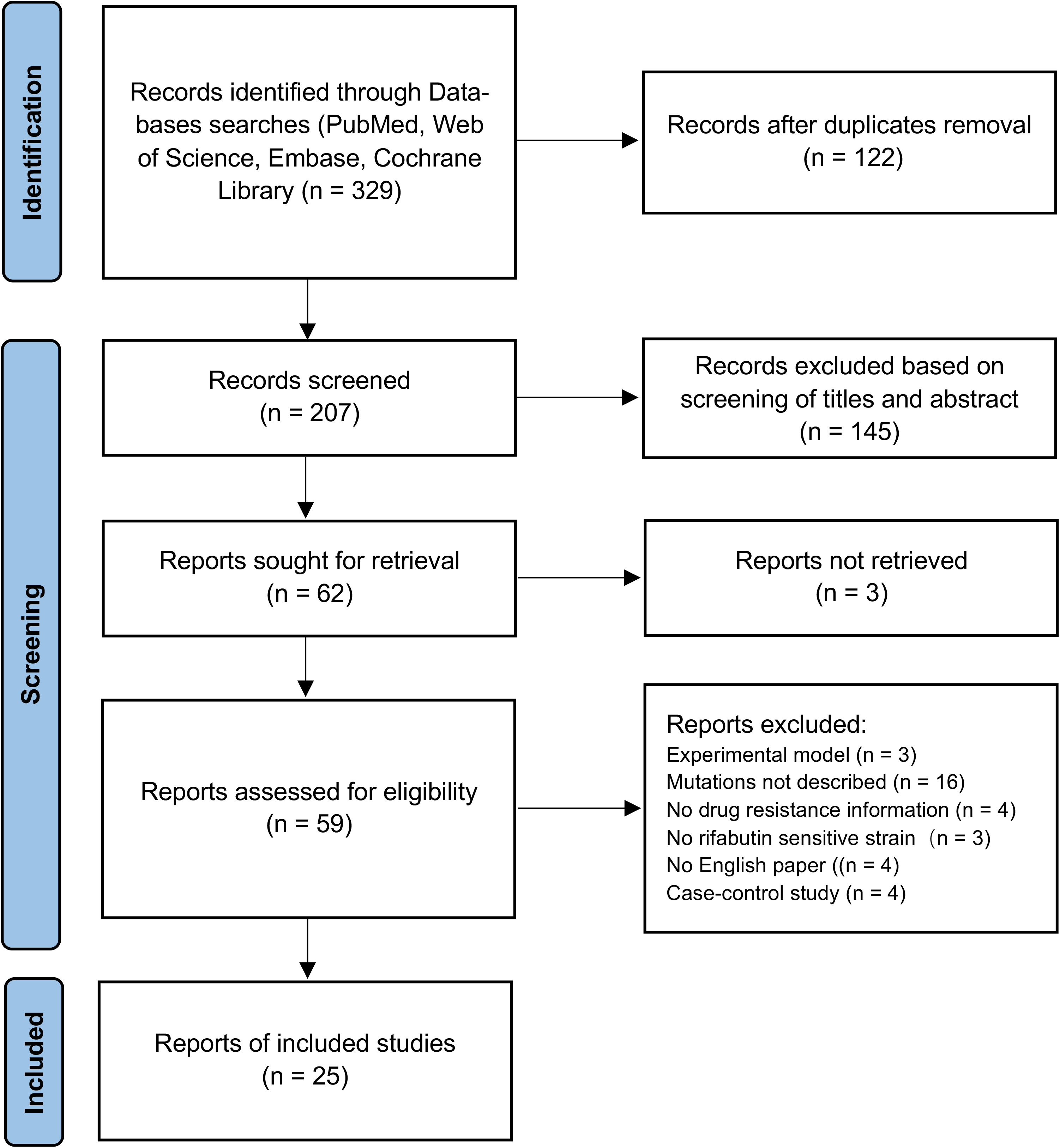
Flow diagram of study selection.

### Quality assessment and characteristics of included studies

To ensure the reliability of the meta-analysis, we first assessed the quality and general characteristics of each included study (Table 1 and Table S2). The results indicated that all the included articles were of low risk of bias and high or moderate quality. pDST for RIF and RFB, were performed using microplate-based assays (11/25), MGIT 960 plate (6/25), BACTEC 460 (4/25), and proportion methods (3/25). There was one study did not provide the pDST method. In 68.00% (17/25) of the articles, the reference methods and standards suggested by the Clinical and Laboratory Standards Institute (CLSI) were employed. Based on the CLSI standards or cutoff specified by the authors, the RIF-resistant isolates in each study were classified as either RFB-susceptible or resistant. Genotypic detection methods primarily involved PCR (16/17) in studies conducted before 2018, while whole-genome sequencing (5/8) became the predominant method from 2019 onwards.

**Table 1.** Characteristics of selected studies (n = 25).

| Author | Study Region | Genotyping method | MIC method | NEW CLSI CBP | Sample size (Experimental/Control) | Quality score |
| --- | --- | --- | --- | --- | --- | --- |
| Yu et al.2022(13) | China | WGS | microplate-based assays | Yes | 6/14 | 6 |
| Li et al.2022(16) | China | WGS | microplate-based assays | Yes | 50/88 | 8 |
| Liu et al.2022(15) | China | WGS | MGIT 960 | Yes | 25/295 | 4 |
| Getahun et al.2022(31) | Ethiopia | MTBDRplus | microplate-based assays | Yes | 9/39 | 8 |
| Li et al.2022(32) | China | PCR | microplate-based assays | No | 18/101 | 5 |
| Nonghanphithak et al.2020(33) | Thailand | WGS | microplate-based assays | Yes | 13/38 | 7 |
| Click et al.2020(34) | Multiple<sup>a</sup> | MTBDRplus | proportion methods | NA | 154/293 | 6 |
| Farhat et al.2019(17) | Peru | WGS | microplate-based assays | Yes | 210/556 | 4 |
| Whitfield et al.2018(21) | Belgium/South Africa | PCR | MGIT 960 | Yes | 27/39 | 6 |
| Jing et al.2017(35) | China | PCR | microplate-based assays | Yes | 52/204 | 8 |
| Rukasha et al.2016(36) | South Africa | PCR | microplate-based assays | Yes | 51/138 | 8 |
| Nosova et al.2016(37) | Russia | TB-TEST | MGIT 960 | Yes | 13/201 | 7 |
| Berrada et al.2016(38) | Multiple<sup>b</sup> | PCR | MGIT 960 | Yes | 31/71 | 6 |
| ElMaraachli et al.2015(39) | South Africa | PCR | MGIT 960 | Yes | 15/42 | 6 |
| Jamieson et al.2014(27) | Canada | PCR | MGIT 960 | Yes | 10/26 | 6 |
| Schön et al.2013(40) | Swedish | PCR | microplate-based assays | Yes | 19/54 | 5 |
| Tan et al.2012(41) | China | PCR | unknown | NA | 43/319 | 5 |
| Chen et al.2012(42) | China | PCR | proportion methods | NA | 101/686 | 6 |
| Yoshida et al.2010(43) | Japan | PCR | microplate-based assays | Yes | 19/23 | 6 |
| Cavusoglu et al.2004(44) | Turkey | PCR | proportion methods | Yes | 6/35 | 6 |
| Yuen et al.1999(45) | Australia | PCR | BACTEC 460 | NA | 6/25 | 6 |
| Sintchenko et al.1999(12) | Australia | PCR | BACTEC 460 | No | 4/17 | 6 |
| Yang et al.1998(11) | Japan | PCR | microplate-based assays | Yes | 15/78 | 5 |
| Williams et al.1998(10) | America | PCR | BACTEC 460 | No | 6/22 | 5 |
| Bodmer et al.1995(9) | Switzerland | PCR | BACTEC 460 | NA | 7/19 | 4 |

Note: a, South Africa, Peru, Philippines, Latvia, Estonia, Thailand, South Korea; b, California, India, Philippines, South Africa, Moldova; CBP, clinical breakpoint; NA, not applicable; Experimental/Control: rifabutin-susceptible isolates/rifabutin-resistant isolates.

These cross-sectional studies included a total of 4333 RIF-resistant clinical MTB isolates from 21 different countries published from 1995 to 2023. Among these isolates, 21.00% (910/4,333) were susceptible to RFB. This proportion varies among different articles, ranging from 6.07% to 45.24% (median: 25.49%). It also varies by country or region (Fig. S1), accounting for 12.80% (42/328) in Europe, 15.61% (348/2,229) in Asia, 19.23% (10/52) in Oceania, 25.00% (16/64) in North America, 26.41% (210/766) in South America, and 28.7% (99/345) in Africa.

### Relationship between RIF resistance mutations and RFB susceptibility

To identify RIF-resistance mutations met the analysis criteria, we counted the mutations reported in these articles. A total of 259 types of *rpoB* mutations were reported (Table S3). Among them, 37 mutations were recommended in the World Health Organization (WHO) catalog (22) as RIF-resistance mutations, of which 28 mutations were reported in two or more studies (median: 4, range: 1-23). The 28 mutations were found in a total of 3,520 (81.23%) RIF-resistant strains. The most common mutation was S450L (2,272 strains, 64.55%), followed by D435V (390 strains, 11.05%), H445Y (211 strains, 5.99%), and H445D (154 strains, 4.38%). Other 24 mutations were listed in Table S4, accounting for 494 strains (14.03%). We subsequently performed a meta-analysis of these 28 mutations.

To investigate the relationship between these mutations and RFB-S, we performed some statistical analyses, including I^2^ statistics, Q-test, and odds ratio (OR) calculation. Based on the results of the I^2^ statistics and Q-test, a random-effects model was used for the meta-analysis of S450L, D435V, and H445C, and a fixed-effects model was used for the analysis of other mutations. The results showed that some mutations at codons 435, 445, 530, 452, and 441 of the *RpoB* were associated with RFB susceptibility (OR>1, P<0.05) (Table 2), including D435V (OR=0.97, 95%CI: 12.32-35.70), D435Y (OR=13.73, 95%CI: 6.83-27.60), D435F (OR=16.04, 95%CI: 5.21-49.43), H445L (OR=18.42, 95%CI: 9.58-35.44), H445C (OR=7.44, 95%CI: 1.62-34.21), H445G (OR=9.82, 95%CI: 2.12-45.53), H445N (OR=9.05, 95%CI: 3.13-26.14), L430P (OR=8.00, 95%CI: 2.13-30.11), L430R (OR=14.43, 95%CI: 2.16-96.32), L452P (OR=6.46, 95%CI: 3.89-10.73), S441L (OR=13.21, 95%CI: 5.53-31.56), and S441Q (OR=11.27, 95%CI: 2.61-48.64).

**Table 2.** Odds ratio of 28 rifampicin resistance mutations occurred in rifabutin-susceptible isolates (experimental) and rifabutin-resistant isolates (control).

| Mutation | No. of studies | No. of isolates | Statistical Method | Effect Estimate | Confidence Level |
| --- | --- | --- | --- | --- | --- |
| D435V | 23 | 389 | OR (M-H, Random, 95% CI) | 20.97 [12.32, 35.70] | High confidence |
| H445L | 17 | 55 | OR (M-H, Fixed, 95% CI) | 18.42 [9.58, 35.44] | High confidence |
| D435F | 7 | 15 | OR (M-H, Fixed, 95% CI) | 16.04 [5.21, 49.43] | High confidence |
| L430R | 2 | 6 | OR (M-H, Fixed, 95% CI) | 14.43 [2.16, 96.32] | High confidence |
| S441L | 13 | 21 | OR (M-H, Fixed, 95% CI) | 13.21 [5.53, 31.56] | High confidence |
| D435Y | 11 | 33 | OR (M-H, Fixed, 95% CI) | 13.73 [6.83, 27.60] | High confidence |
| S441Q | 3 | 5 | OR (M-H, Fixed, 95% CI) | 11.27 [2.61, 48.64] | High confidence |
| H445G | 4 | 5 | OR (M-H, Fixed, 95% CI) | 9.82 [2.12, 45.53] | Moderate confidence |
| H445N | 4 | 15 | OR (M-H, Fixed, 95% CI) | 9.05 [3.13, 26.14] | Moderate confidence |
| L430P | 5 | 6 | OR (M-H, Fixed, 95% CI) | 8.00 [2.13, 30.11] | Moderate confidence |
| H445C | 8 | 23 | OR (M-H, Random, 95% CI) | 7.44 [1.62, 34.21] | Moderate confidence |
| L452P | 11 | 62 | OR (M-H, Fixed, 95% CI) | 6.46 [3.89, 10.73] | Moderate confidence |
| I491F | 3 | 9 | OR (M-H, Fixed, 95% CI) | 2.72 [0.80, 9.30] | Indeterminate |
| S450Q | 3 | 7 | OR (M-H, Fixed, 95% CI) | 2.27 [0.61, 8.37] | Indeterminate |
| H445S | 6 | 7 | OR (M-H, Fixed, 95% CI) | 2.10 [0.70, 6.35] | Indeterminate |
| H445Q | 3 | 6 | OR (M-H, Fixed, 95% CI) | 1.00 [0.16, 6.07] | Indeterminate |
| Q432P | 8 | 20 | OR (M-H, Fixed, 95% CI) | 0.95 [0.36, 2.49] | Indeterminate |
| V170F | 5 | 12 | OR (M-H, Fixed, 95% CI) | 0.93 [0.23, 3.70] | Indeterminate |
| Q432E | 3 | 3 | OR (M-H, Fixed, 95% CI) | 0.81 [0.09, 7.16] | Indeterminate |
| H445P | 7 | 10 | OR (M-H, Fixed, 95% CI) | 0.79 [0.24, 2.58] | Indeterminate |
| Q432L | 7 | 13 | OR (M-H, Fixed, 95% CI) | 0.68 [0.21, 2.22] | Indeterminate |
| S450F | 5 | 6 | OR (M-H, Fixed, 95% CI) | 0.66 [0.16, 2.74] | Indeterminate |
| H445R | 16 | 71 | OR (M-H, Fixed, 95% CI) | 0.52 [0.25, 1.06] | Indeterminate |
| S450W | 13 | 56 | OR (M-H, Fixed, 95% CI) | 0.46 [0.22, 0.98] | No association |
| H445Y | 21 | 211 | OR (M-H, Fixed, 95% CI) | 0.43 [0.27, 0.68] | No association |
| H445D | 20 | 154 | OR (M-H, Fixed, 95% CI) | 0.38 [0.22, 0.65] | No association |
| Q432K | 7 | 28 | OR (M-H, Fixed, 95% CI) | 0.33 [0.11, 1.02] | Indeterminate |
| S450L | 23 | 2272 | OR (M-H, Random, 95% CI) | 0.09 [0.05, 0.16] | No association |

Abbreviations: No, number; OR, Odds ratio; CI, confidence interval.

These mutations were further evaluated for their confidence grade in the prediction of RFB-susceptibility according to the WHO guidelines (23). We found that seven mutations, namely D435V, H445L, D435F, L430R, S441L, D435Y, and S441Q, had high confidence (Fig. 2). Strains carrying these mutations showed a rate of 83% (435/524) susceptibility to RFB. Among these mutations, the most prevalent one was D435V (34.18%, 311/910), followed by H445L (5.60%, 51/910) and D435Y (3.19%, 29/910). Five mutations, H445C, H445G, H445N, L430P, and L452P, were moderate confidence in predicting RFB-susceptibility (Fig. S2), with 62.16% (69/111) of the strains carrying these mutations susceptible to RFB.

**Fig 2.**
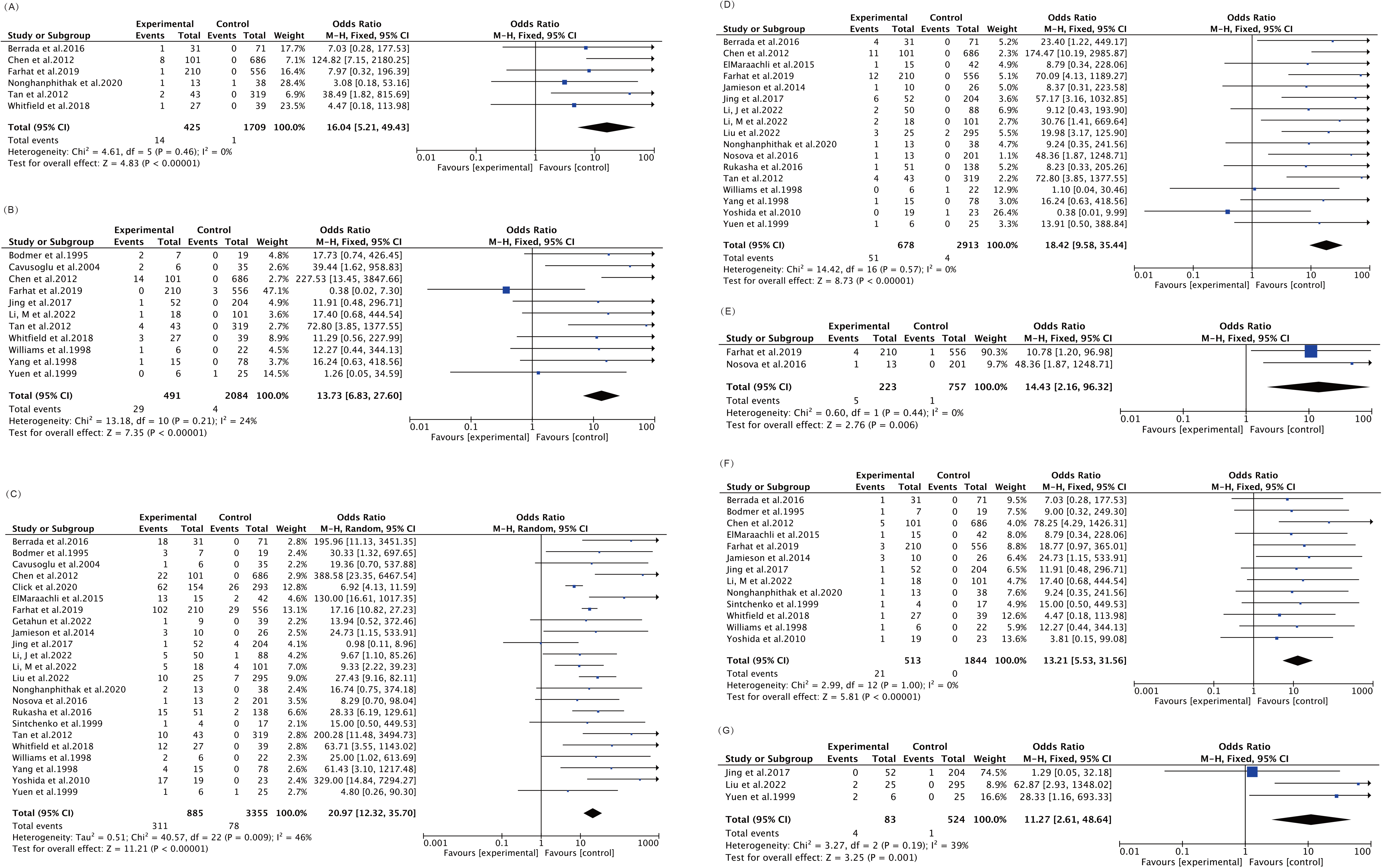
Forest plot for odds ratio of rifampicin resistance mutations D435F(A), D435Y(B), D435V (C), H445L(D), L430R(E), S441L(F), S441Q(G) occurred in rifabutin-susceptible isolates (experimental) and rifabutin-resistant isolates (control).

We also found that four mutations, S450L (OR=0.09, 95% CI:0.05-0.1), S450W (OR=0.46, 95% CI:0.22-0.98), H445Y (OR=0.43, 95% CI: 0.27-0.68), and H445D (OR=0.38, 95% CI: 0.22-0.6), were significantly associated with RFB-resistance phenotype (OR<1) (Table 2). Other mutations did not show a significant correlation with RFB susceptible or resistant phenotype.

### Heterogeneity, susceptibility, and publication bias analysis

To assess the reliability of the aforementioned analysis results, we conducted homogeneity, sensitivity, and bias analyses for the included studies and mutations. Heterogeneity was observed for S450L (I^2^=74%, P_het_<0.00001), D435V (I^2^=46%, P_het_=0.009), and H445C (I^2^=49%, P_het_=0.06). However, meta-regression analyses based on publication year, study location, sample size, drug resistance detection method, and mutation detection method failed to explain the heterogeneity (Table S5). The remaining mutations showed low heterogeneity (Table S4).

In the sensitivity analysis of the results, the influence of each individual data set on the pooled OR was assessed by removing each study individually. No obvious changes were found for all mutations except for the H445Y mutation. Excluding the results from certain studies, the combined results were not robust (95%CI including 1), indicating that those results for H445Y were not reliable (Fig. S3).

The results of Egger’s test (*p*=0.007) and funnel plot (asymmetry) revealed that publication bias has an impact on the results of S441L (Table S4). However, no discernible changes were found in the findings when excluding Chen et al.’s study (Fig. S4), indicating there was no impact of this publishing bias on the overall findings.

### Additional analysis

The relationship between drug resistance mutations and the resistance levels of the corresponding drug in MTB is of significant importance for clinical TB treatment (24-26). Therefore, we further investigated the minimum inhibitory concentrations (MICs) of strains with RFB susceptibility-associated mutations to RFB and RIF. A total of 1842 strains collected from 15 studies that used the CLSI criteria for DST and provided MICs for both drugs were included in this analysis. Strains carrying the high-confidence mutations showed low MIC (D435V, D435F, H445L, and D435Y) or were fully susceptible (S441L) to RFB (Fig. 3A), but had high (D435V and D435F) or moderate (H445L, D435Y, and S441L) MICs to RIF (Fig. S5). Moreover, the MIC distributions of these mutant strains for both drugs were significantly unrelated (R^2^=0.096, p-value<0.001) (Fig. 3B). Only one MTB strain carrying the L430R and S441Q mutations had MIC result, and additional data were needed to establish the two mutations with resistance levels of RFB and RIF.

**Fig 3.**
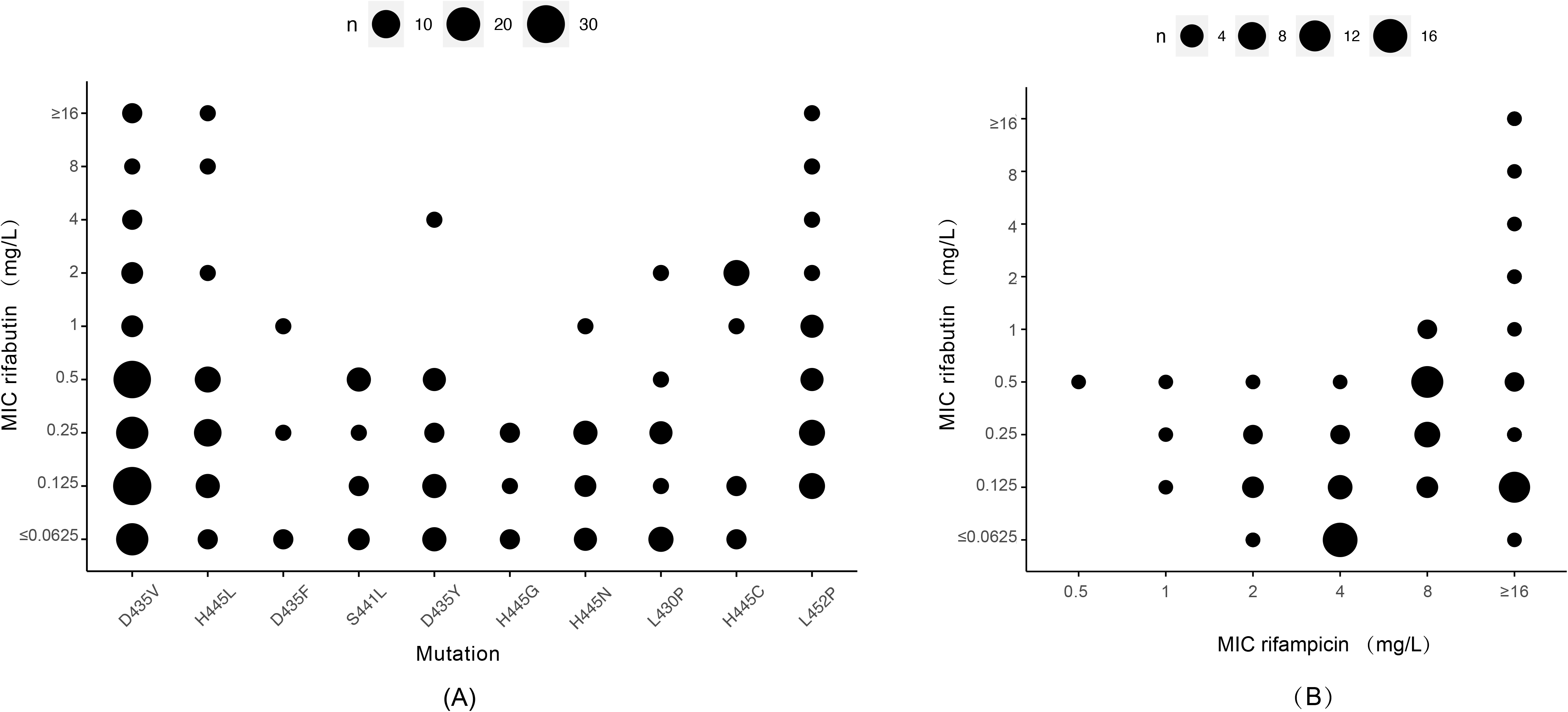
The minimum inhibition concentrations (MIC) of isolates carrying mutations with high confidence in prediction of rifabutin-susceptibility to rifabutin (A) and to both rifampicin and rifabutin (B).

Strains carrying mutations with moderate confidence showed moderate (H445C, L452P), low (H445N, L430P) MICs, or susceptibility (H445G) to RFB, but had high (H445C) or moderate (L452P, H445G, H445N, L430P) MICs to RIF (Fig.S5).

## Discussion

This study, by compiling 25 studies, obtained *rpoB* mutations and pDST results of RFB for 4,333 clinical RIF-resistant MTB isolates from 21 countries. Among these RIF-resistant strains, 20.9% (910/4333) were found to be susceptible to RFB. Certain RIF-resistance mutations on *rpoB* codons 435, 445, 441, 430, and 452 were associated with the RFB-susceptible phenotype, including seven high confidence mutations (D435V, D435F, H445L, D435Y, S441L, L430R, S441Q) and five moderate confidence mutations (H445C, H445G, H445N, L430P, L452P). Among RR-TB strains carrying these mutations, 83.01% and 62.16% were susceptible to RFB. In contrast, *rpoB* mutations H445D, H445Y, S450L, and S450W were associated with RFB-resistance.

Notably, several investigations revealed that partial RIF-resistant strains were susceptible to RFB, but the proportions varied. We found in this meta-analysis that the proportion was about 20.9%, suggesting that a considerable proportion of MDR/RR-TB patients may benefit from treatment with RFB. However, this proportion varies across different regions, higher in Africa and America and lower in Asia and Europe, which may be attributed to variations in the types of mutations carried by local strains. Previous studies have indicated that the genetic background of strains also influence the level of resistance of strains with same mutation. For example, Farhat et al (17) had observed that Beijing strains carrying the D435V mutation had a higher MIC to RFB than Lineage 4 strains in a clinical MTB isolate library. Further research can combine the influence of genetic background and interaction of mutations to provide more references for the use of RFB.

Although many studies have explored the association of certain RIF-resistance mutations with RFB-susceptibility, the limited number of mutant strains has hindered a more reliable assessment of the results. Meta-analysis overcomes this limitation by consolidating data from multiple studies. We found that *rpoB* D435V (*E. coli* D516V) was the most significant mutation associated with RFB susceptibility, with a frequency of 34.18% among RFB-susceptible strains. Although there was heterogeneity, most relevant studies (22/25) supported this association (Fig.2C). Furthermore, Gill, S. K. et al. have confirmed this correlation through the construction of site-directed mutant strains (18). The frequencies of other mutations were much lower than D435V, and their correlation with RFB susceptibility had not been previously evaluated. However, Anthony et al (19) found that S441L was associated with RFB susceptibility in an *in vitro* assay.

The low MIC levels observed in corresponding mutant strains further validate the reliability of the high confidence mutations we assessed for predicting RFB susceptibility. The significant lack of correlation between MICs for RFB and RIF further supports the potential use of RFB as an alternative treatment for MDR/RR-TB caused by these strains. However, whether RFB can effectively treat such TB patients needs to be determined through retrospective analysis of corresponding clinical treatment outcomes.

Interestingly, we found that most of different mutations in *rpoB* gene at the same codon were only associated with either RFB susceptibility or resistance, such as those at codons 435, 441, 448 and 450. However, certain mutations at specific codons were associated with both phenotypes, such as mutations at codon 445 (E. coli 526). H445L, H445C, H445G, and H445N were associated with RFB susceptibility, while H445Y and H445D were associated with resistance. This was similar to the finding by Jamieson *et al.* (27) that different mutations at codon 445 were associated with different levels of drug resistance or susceptibility of RIF, which may be due to differences in spatial configurations caused by different mutations (28). This indicates that the development of new diagnostics requires the detection of specific mutations, not just mutation sites.

This study had certain limitations. Firstly, the relationships between nine of the 37 known RIF-resistance mutations were not estimated, due to they were only reported in one study. Secondly, the effect of genetic background on the resistance level were not analyzed because of the absence of such information in many isolates. Thirdly, there were some missing data in susceptibility results and potential biases in the included studies. These limitations have little effect on the reliability of the mutations we identified as being associated with RFB susceptibility.

In conclusion, through meta-analysis, we have elucidated the proportion of RFB susceptibility among RIF-resistant strains and identified the RIF-resistance mutations associated with such phenotype. We have demonstrated the high accuracy of predicting RFB-susceptibility using mutations such as *rpoB* D435V, R448L, H445L, D435F, L430R, S441L, D435Y, S441Q, and D435E/S441L. This provides a theoretical basis for the development of rapid detection methods for clinical RFB-susceptible strains and the use of RFB in treating MDR/RR-TB.

## Materials and methods

### Search strategy and selection criteria

A comprehensive literature search was conducted in the PubMed, Web of Science, Embase, and Cochrane Library databases until June 1, 2023. The search terms included *Mycobacterium tuberculosis*, rifabutin, rifampicin, resistance, mutations, and their variants, with no language or date restrictions (Table S1). Potentially relevant studies were retrieved and reviewed by two reviewers. All records from different sources included in our search strategy were merged and uploaded to the reference management software Endnote, and duplicates were removed. We included cross-sectional studies that involved *rpoB* mutations and pDST results for RIF and RFB. Only full-text English-language articles were considered. Literatures such as reviews, conference abstracts, comments, and news articles were excluded. Studies that focused on experimental strains, without RFB-resistant or susceptible groups, and did not provide detailed information on mutations or pDST results were also excluded. The PRISMA flow diagram (29) was used to illustrate this process, and the reasons for excluding studies were documented in detail (Fig. 1).

### Data extraction

Two reviewers (W.W. and T.Y.) independently used standardized data extraction spreadsheets to collect information and assess the quality of the included studies. If an article included various anti-tuberculosis drugs, we only extracted data pertaining to RIF and RFB. The data we extracted were include author(s), publication year, study period, study location, sample size, pDST and MIC to RIF, pDST and MIC to RFB, mutations and the detection methods, and methods for pDST/MIC.

### Quality assessment

The included cross-sectional studies were evaluated using the bias risk assessment criteria recommended by the Agency for Healthcare Research and Quality (AHRQ). The AHRQ consists of 11 items, and each item in the AHRQ was answered with "yes (one score)", "no (zero score)" or "not reported (zero score)". A score of 8-11 indicates high quality, 4-6 indicates moderate quality, and 0-3 indicates low quality. Only studies with low bias risk and high or moderate quality (no high bias risk) were included in meta-analysis.

### Sources of drug-resistant mutations and Statistical analysis

The RIF-resistance mutations were sourced from the 2021 WHO’s catalogs (22), which includes mutations with a Final Confidence Grading of “Assocw R” and “Assocw RI”. Meta-analyses were carried out with RevMan (version 5.4) and Stata 16.0 software. The odds ratios (OR) were estimated with a 95% confidence interval (CI). The OR>1, OR<1, and OR=1 indicated the mutation was correlated with RFB susceptibility, correlation with RFB resistance, and no correlation found between it with both phenotypes, respectively. Additionally, we evaluated the confidence grade of each mutation in the prediction of RFB susceptibility using the method applied in the WHO’s 2018 technical guidelines (23). Mutations were categorized into five confidence levels based on thresholds, including High confidence (OR>10, *P*<0.05), Moderate confidence (5< OR≤10, *P* <0.05), Minimal confidence (1<OR≤ 5, *P*<0.05), No association (OR < 1, *P* < 0.05), or Indeterminate (*P*≥0.05).

Heterogeneity analysis was assessed using the Cochrane Q test. When P_het_ > 0.1 and I^2^ ≤ 50%, there was no significant heterogeneity among the studies, and a fixed-effect model was used for meta-analysis. When P_het_ ≤ 0.1 or I^2^ > 50%, a random-effects model was applied. If there were fewer than five included studies, a fixed-effect model was performed. In cases of high heterogeneity, meta-regression analysis was conducted to explore potential associations between publication year, study location, sample size, drug resistance detection method, mutation detection method, and study results (i.e., mutation and RFB susceptibility). Sensitivity analysis was performed using the one-by-one exclusion method to evaluate the stability of the results. Publication bias were assessed by Funnel plots and Egger’s test.

### Additional analysis

To assess the relationship between RFB susceptibility-associated mutations and MIC levels to RFB, we conducted additional analysis after the meta-analysis. The MIC levels of each mutation were evaluated according to the criteria set by WHO (23). Based on whether the determined MICs crossed the Critical Concentration (CC), they were categorized as follows: High MICs, none of the identified/available MIC distributions (in any media) span the current CC; Moderate MICs, some of the identified/available MIC distributions (in any media) span the current CC; Low MICs, the majority of identified/available MIC distributions, across different media, included the current CC; Need Additional Data, conflicting MIC results for any media or too few data points to establish conferred resistance (e.g. Only 1 isolate tested in any media). According to the standards of the CLSI M24-A2(30), the CC for RIF and RFB were 1 μg/ml and 0.5 μg/ml, respectively.

## Acknowledgments

This study was supported by project of Health Commission of Zhejiang Province (2023RC065, 2023RC066), and Hangzhou Agricultural and Social Development Scientific Research Project (202204B02).

